# Alpha-synuclein overexpression reduces neural activity within a basal ganglia vocal nucleus in a zebra finch model

**DOI:** 10.1101/2025.09.11.675693

**Authors:** Brian R. Dominguez, Gabriel Holguin, Madeleine S. Daly, Reed T. Bjork, Stephen L. Cowen, Julie E. Miller

## Abstract

Changes in vocal pitch, loudness, and timing are prevalent in Parkinson’s Disease (PD) and a target for early intervention and treatment. The neural mechanisms underlying these impairments are not understood, motivating work in animal models. The adult male zebra finch songbird is uniquely poised for these studies given vocally-dedicated brain nuclei and a quantifiable output (birdsong). Our prior publication revealed that injection of an adeno-associated virus (AAV5) expressing the human (h) alpha-synuclein (h*SNCA*, a-syn) gene into basal ganglia vocal nucleus Area X results in elevated insoluble a-syn protein and parkinsonian-like changes including softer, shorter, and reduced vocalizations compared to controls. Here, we test the hypothesis that AAV-h*SNCA* overexpression reduces the firing rate of specific neuronal sub-types in Area X using *in vivo* recordings in anesthetized finches. Five classes of neurons were differentiated in AAV-treated finches based on waveform width (narrow vs. wide) and firing rates (low vs. fast). Wide-waveform/low-rate activity is a consistent feature of striatal medium spiny neurons (MSNs), a dominant cell type in Area X and in mammalian basal ganglia. Reduced firing rates and enhanced post-peak rebound were detected in the AAV-h*SNCA* group for putative MSN neurons compared to AAV controls. No differences in firing rate nor waveform shape were detected for the narrow waveform neurons. Our findings provide the first characterization of early a-syn-driven neural activity changes in vocal control neurocircuitry.

## Introduction

Pathological accumulation of alpha-synuclein (a-syn) is believed to contribute to multiple neurodegenerative diseases, including Parkinson’s disease (PD). Accumulation of toxic a-syn species culminates in the development of Lewy Bodies and Lewy Neurites that are detected in inherited and sporadic cases of PD, dementia with Lewy Body disease, and Multiple Systems Atrophy (1–5). How abnormal increases in a-syn protein expression and its aggregation disrupts ongoing neuronal activity in the basal ganglia (BG) is not understood, yet this information is critical to developing cell-specific therapeutic targets (6–8). One experimental approach is to examine how elevated a-syn protein interferes with the ongoing firing activity of striatal neurons.

Elevated levels of a-syn protein at cortico-striatal synapses can result in synaptic failure and loss of cortico-striatal plasticity based on electrophysiological recordings from rodent brain slices (6–11). Striatal Medium Spiny Neurons (MSNs) show alterations in dendritic spine density, interactions between a-syn aggregates and their NMDA receptors, and reduced efficacy of cortical pre-synaptic glutamatergic input that affects their firing rate (12–15). Furthermore, burst firing is a characteristic feature of MSNs (16), and dysregulated MSN burst firing may contribute to PD as shown in 6-hydroxydopamine (6-OHDA) models (17). A recent study used induced human pluripotent stem cells from patients with Multiple System Atrophy of the Parkinsonian type that were differentiated into MSNs. These MSNs showed reduced spontaneous firing rates, frequency, and amplitude of miniature post-synaptic currents. These changes were also associated with increased a-syn release (18). How *in vivo* cell-specific neuronal firing patterns in the basal ganglia relates to the behavioral output is unclear and a critical gap. The scant literature on *in vivo* recordings in rodent h*SNCA* overexpression models comes from nigral, thalamic, and hippocampal neurons evaluating movement and cognition (19–21). Pairing electrophysiological changes in basal ganglia circuitry with vocal behavior offers an early and critical entry-point into furthering our understanding of the effects of alpha-synucleinopathies on neural circuits.

Voice and speech deficits are prevalent in PD and affect quality of life (22, 23). Early stage vocal deficits (e.g. a soft, monotonous voice, altered speaking rate) can appear years before the movement symptoms, highlighting their utility as an early diagnostic tool. The use of machine learning algorithms and smartphone applications also support using vocal changes as an early biomarker of PD (24–37). Our group and others have modeled PD-related vocal deficits using pharmacological and genetic approaches in rodents and songbirds with data from these experiments implicating the h*SNCA* gene in driving early vocal dysfunction (38–46).

Songbird models offer distinct advantages over rodents in exploring how the brain encodes sensory and motor information, shaping vocal learning and production in response to feedback (47). Males of the zebra finch species sing but the females do not. The male finch song control system in the brain **(Fig 1A**) has similar neural organization, genetics, and physiology of the basal ganglia to mammalian species (48–50). The song control system has strong similarities to human speech and auditory centers in the brain (51–57). Vocal learning and on-going song modification in the zebra finch model can be investigated through the anterior forebrain pathway, a structure homologous to the cortico-basal ganglia-thalamo-cortical loop in humans and other mammals (50). Song-dedicated brain region Area X in the basal ganglia shares similar genetics to human striatum, a major area involved in speech initiation and learning, and has cell types characteristic of mammalian neuronal populations (48, 54, 58, 59).

**Fig 1.**
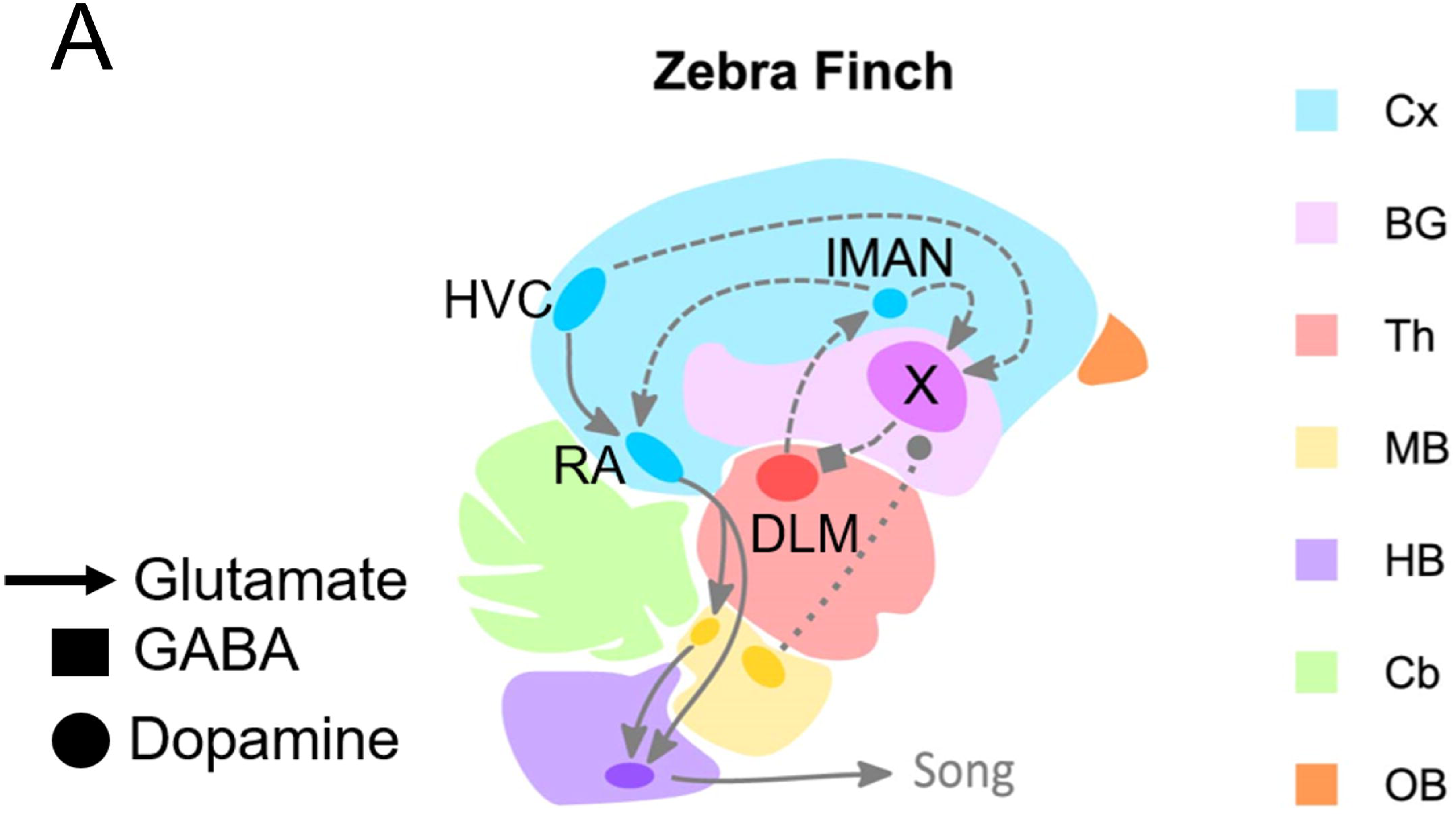
Finch song control circuitry. Brain nuclei in the adult male finch are specialized for song learning and production (HVC and Area X: proper names; RA: robust nucleus of the Arcopallium, lMAN: lateral magnocellular nucleus of the anterior nidopallium). The solid lines represent cortico-brainstem circuits involved in vocal production (finch: HVC to RA to brainstem). The dashed lines represent cortico-basal ganglia-thalamo-cortico circuits for on-going vocal learning and modification (lMAN-Area X-DLM-lMAN). Abbreviations: Cx – cortex. BG – basal ganglia. DLM – dorsal medial nucleus of the thalamus. Th – thalamus. MB – midbrain. HB – hindbrain. Cb – cerebellum. OB – olfactory bulb. Area X (violet) contains striatal, globus pallidus, and interneurons but in the human BG, striatal and pallidal areas are separate. Several auditory areas provide input to these vocal nuclei not shown here) (60, 61). Figure and legend modified from finch brain image in Fig 1 of Medina et al. (2022).

Electrophysiological recordings in singing finches and anatomical tracings show that MSNs are the dominant cell type in Area X and drive firing activity of globus pallidus-like (PAL) projection neurons (50, 58, 62). As in mammals, GABAergic MSNs and Globus-pallidus (GP, PAL) type neurons receive glutamatergic (Glu) inputs from cortex (in finch-song nuclei HVC and lMAN, **Fig 1A**). GP neurons project to the dorsal medial thalamic nucleus (DLM) which then relays its output to the cortex (63). While studies of MSN activity in animal models and humans are difficult due to low spontaneous activity (14, 64), finch MSNs are dynamically active during singing and show temporally precise relationships to song structure (62, 65). DLM projections to HVC can also modulate syllable transitions (66). High tonic firing in these PAL neurons inhibits DLM, reducing excitation onto lMAN and then subsequently onto song nucleus RA, leading to altered song output including variations in pitch (55, 62). Area X and lMAN neurons exhibit firing variability that is related to birdsong features (55, 62). The well-characterized neuronal populations in these brain nuclei and circuitry connections make this a powerful model to study the impact of alpha-synucleinopathies on vocalizations.

In our prior work, we showed that targeting Area X with an adeno-associated virus (AAV) driving increased expression of the human (h) wild-type *SNCA* gene leads to PD-like changes in the song including reduced duration and intensity. (45). We also showed that a-syn overexpression results in protein aggregates in Area X cell bodies (46). Here, as a critical first step, we categorize cell-specific firing patterns and waveforms in Area X that are affected by a-syn overexpression compared to controls using tetrode recordings in anesthetized finches immediately following singing behavior. Our working hypothesis (**Fig. 1B**) is that h*SNCA* overexpression in Area X will decrease firing rates and alter burst-firing in putative Area X MSNs. We tested this hypothesis by recording from Area X neurons in anesthetized zebra finches.

**Fig 1B.**
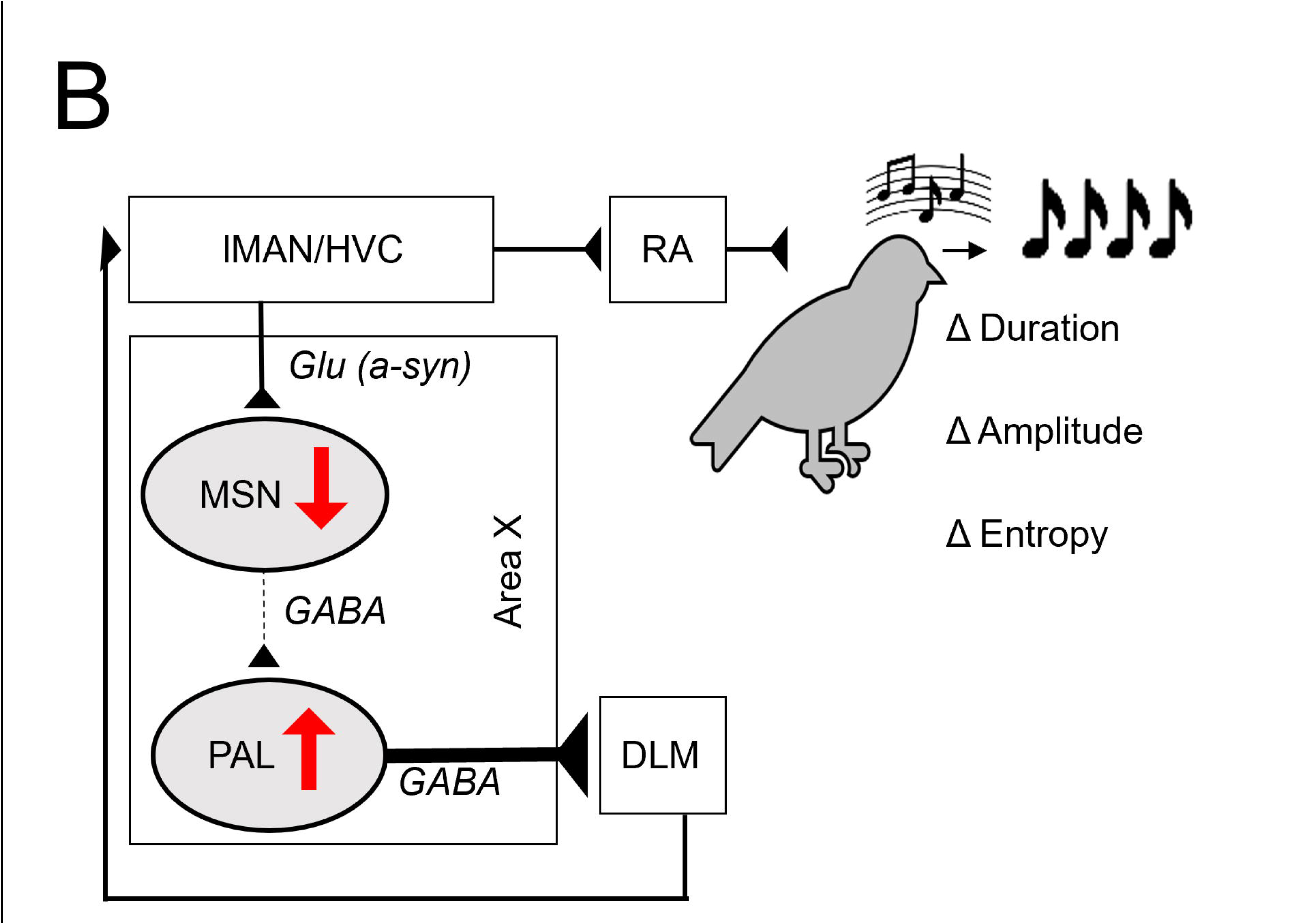
Hypothetical model for a-syn driven changes in neural activity in Area X and consequences on song control circuitry. Cortical vocal nuclei (lMAN, HVC) provide glutamatergic (Glu) input to Area X neurons. Striatal Area X MSNs provide GABAergic input to globus-pallidus like (GP, PAL) neurons including those that inhibit thalamic nucleus DLM. Here, we *predict* that abnormally elevated levels of AAV-driven a-syn protein in Area X could alter intrinsic membrane properties of MSNs and/or reduce Glu input to MSNs. Consequently, MSNs would reduce their firing rate (red arrow), leading to increased GP PAL projection neuron firing and increased inhibition of DLM. Increased DLM inhibition could be responsible for changes in amplitude (loudness) and duration of the song syllables in finches with the AAV-h*SNCA* phenotype (45).

## Materials and Methods

### 2.1. Finch subjects, song recording and analyses

All animal use was approved by the Institutional Care and Use Committee at the University of Arizona. Details on finch numbers, criteria for study inclusion/exclusion, and experimental endpoints are provided in the following sub-sections. **Fig 1C** shows our experimental timeline, workflow, and tetrode array.

**Fig 1C.**
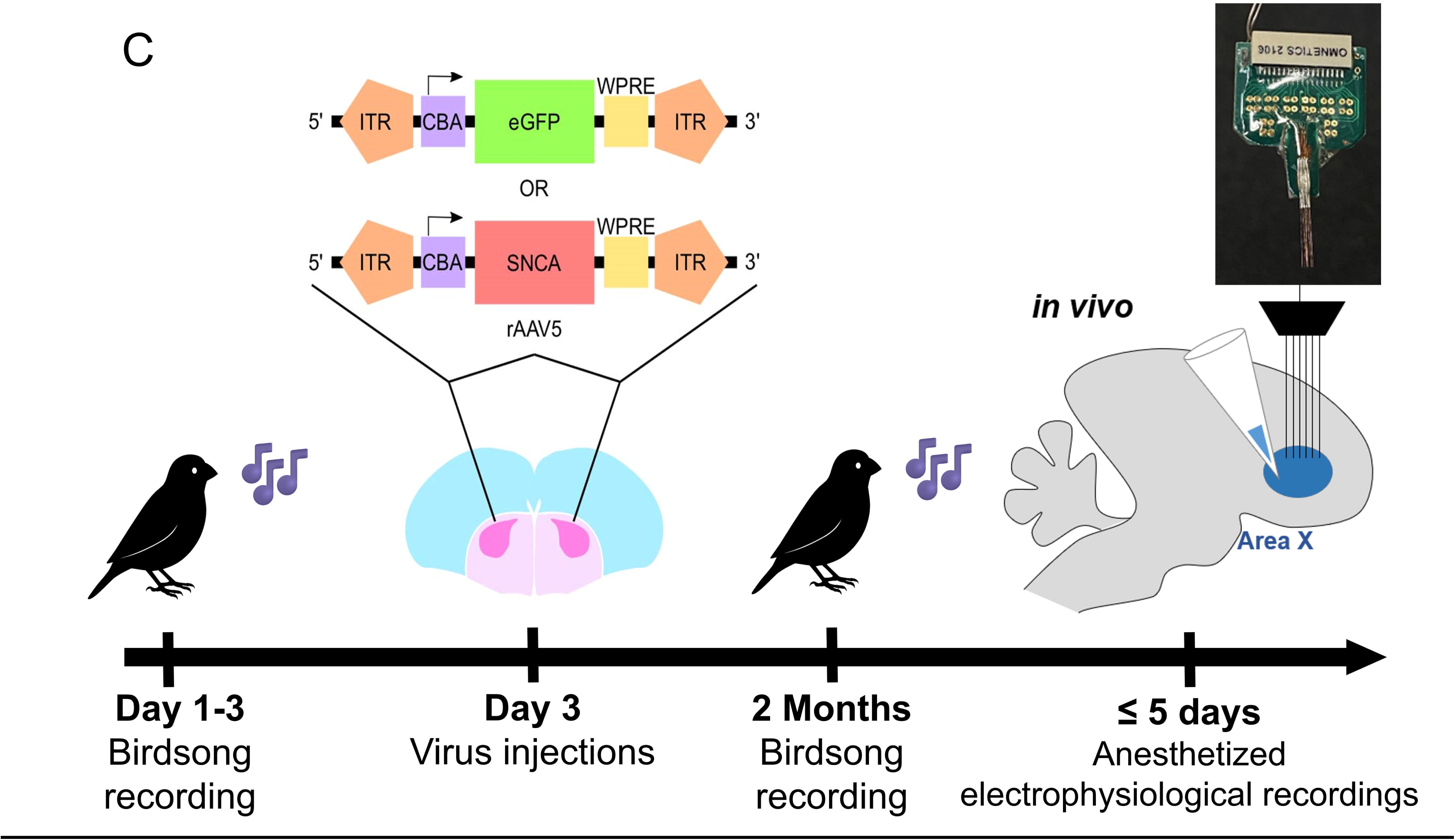
Tetrode array and experimental timeline. Schematic of AAV constructs and coronal section showing bilateral song nucleus Area X (purple) modified from Fig 3b in Medina et al. (2022). Song was collected within three days of the bilateral injection into Area X and then again at two months post-virus injection followed by tetrode recordings in anesthetized finches within five days, for a total experimental duration of approximately two and a half months. At the end of the recording morning (≤3 hrs), the finch was humanely euthanized, and the brain removed post-mortem for cryosectioning followed by immunohistochemistry to confirm AAV and a-syn protein expression along with tetrode placement.

Animal procedures followed Medina et al. (2022) and are summarized here. Only finches that showed normal eating/drinking, perching, and movement in their cages were selected for the study. Finches that did not sing prior to the pre-injection day were excluded from the study and returned to the general aviary population. Adult non-breeding male zebra finches (approx. 197-365 days post-hatch at time of euthanasia) were moved into individual sound attenuation chambers (Eckel Noise Control Technologies) and acclimated under a 14:10 h light-dark cycle for at least two days prior to song recordings and neurosurgery. Behavioral experiments were conducted in the morning from lights-on for two hours to match our prior study (45) along with ensuring singing-driven neuronal activity just prior to conducting the tetrode recordings in anesthetized finches.

#### Song recordings and analyses

Songs were acquired from individual males singing alone (known as undirected song) using free-field omnidirectional microphones (Shure 93) suspended at set points over the center of the cage, connected to an audiobox (Audiobox: 44.1 kHz sampling rate per 24-bit depth; Niles, IL) and recorded with Sound Analysis Pro 2011 (SAP 2011, (67) http://soundanalysispro.com/). Song data collected and used for this study can be found at the University of Arizona ReData repository upon publication (10.25422/azu.data.27868266).

A basic unit of birdsong is defined as a motif; a motif is comprised of a sequence of repeated syllables (**S1 Fig A**). Wav files from the two-hour recording period were joined (Shuang’s audio joiner) and then the entire file was viewed in Audacity (https://www.audacityteam.org/). The most frequently occurring motif was then identified in the joined wav file for two pre-surgical mornings (time point 0) and then two post-surgical mornings (time point, 2 months, **Fig 1C**) by an investigator blind to the AAV condition. The two-month timepoint was selected as the experimental endpoint for song analyses and tetrode recordings in a non-survival surgery followed by humane euthanasia. Our prior publications show that the most prominent song and protein changes occurred at this time point in the AAV-h*SNCA* finches (45, 46). Following morning lights-on, the first motif was identified in the wav file and then 25 consecutive renditions of the individual syllable wav files were exported from Audacity from the four mornings. Up to five unique syllables within a motif for each bird were included in the analyses.

Syllable wav files were processed in Matlab R2014 using the SAP SAT Tools box using code (provided by Dr. Nancy Day, Whitman College) to extract individual acoustic features as a feature batch function. Raw amplitude (intensity) measurements were calibrated based on sound chamber location as in our prior study (68). Mean, standard deviation, and coefficient of variation (CV, standard deviation/mean) were then calculated by averaging 50 scores (25 syllable copies x 2 recording sessions) across time points 0 (pre-AAV injection) and 2 months post-AAV injection for each syllable within the bird’s motifs to account for natural, day to day fluctuations in song. For group level comparisons, a normalization score was obtained per bird by dividing timepoint two months (2) post-injection by pre-injection (0). After verifying that scores were not normally distributed, scores were then compared using the Mann Whitney U test in GraphPad Prism (version 10) between AAV-h*SNCA* and AAV-GFP groups for all syllable types combined and then sorted by syllable type (**S1 Table**) given prior findings on a-syn differential effects (45). Syllables were sorted into noisy, mixed, or harmonic categories based on visual inspection and Wiener Entropy (WE) scores following our prior publications (**S1 Fig**) (45, 69). We focused our analyses on comparing duration, amplitude, and WE scores between the two AAV groups, and their correlations with the electrophysiological analyses for an individual bird (see section on **Statistics** and **S1-2 Figs**). A subset of these birds was used in our separate publication evaluating regional distribution of a-syn protein in finch song nuclei and linear correlations to syllable features in Area X and lMAN (46).

As described in Medina et al. (2022), duration is the length of each syllable (in seconds), and amplitude is the intensity or loudness of the syllable (in decibels-dB). WE is a measure of uniformity in the power spectra (i.e., structure of the syllable), where scores are calculated on a logarithmic scale; noisy syllables have values approaching zero while harmonic syllables have more negative scores. The coefficient of variation (CV) is calculated by dividing the standard deviation of 50 syllable renditions by the mean score and represents variability across motifs for an acoustic feature, modulated by the lMAN-Area X-DLM-lMAN loop (70).

### 2.2 Surgical procedures: viral injections

A total of 19 finches were originally used for adeno-associated virus (AAV) bilateral injections into Area X (n=11, AAV5-CBA-h*SNCA* [human alpha-synuclein]; n=8, AAV5-CBA-eGFP controls [Green Fluorescent Protein]; CBA: chicken beta-actin promoter). One bird in each group spontaneously died prior to the two month post-AAV injection experimental endpoint due to unknown causes and therefore, they were removed from the study. We alternated the order of the AAV experimental vs. control virus surgeries when two surgeries were done on the same day.

The surgery suite and instruments were prepared for aseptic survival surgery under PI Miller’s approved IACUC Protocol 13-489. Following two hours of morning song collection, finches were put under isoflurane anesthesia with oxygenation (2-3%, 0.5 liters/min oxygen) and set up in the stereotaxis. The analgesic lidocaine (0.05ml) was subcutaneously injected across the planned incision site of the scalp at four locations. For the bilateral injections into Area X, stereotaxic coordinates were used from the bifurcation of the mid-sagittal sinus: 3.1 mm rostral-caudal, 1.62 mm medio-lateral, a depth of 3.1 mm, and head angle of 40 degrees. These coordinates avoid indirect targeting of song nucleus lMAN with virus as previously described (45). Viruses (AAV5) were obtained from the University of North Carolina Viral Vector Core through financial support of the Michael J. Fox Foundation for Parkinson’s Research (https://www.michaeljfox.org/research-tools).

A glass pipette was fitted into a Nanoject II pressure-injector and back-filled with mineral oil, then loaded with either AAV. Approximately 500 nL of virus was injected at a rate of 27.6 nL/injection every 15 seconds for a total of 18 injections followed by five minutes of wait time then inspection of pipette tip for clogging. Upon removal of isoflurane anesthesia, birds were monitored for any signs of pain and distress while on a homeothermic heating blanket. Within a half-hour, birds were noted to be alert, upright, and vocalizing and did not require any post-operative analgesia. They were returned to their sound chambers and monitored for any surgical complications, abnormal postural or movement, eating and drinking behavior for at least three days post-surgery by research personnel fully trained in bird care. Their drinking water was supplemented with an oral antibiotic dose of TMS (Sulfamethoxazole and Trimethoprim Oral Suspension, USP grade; 1ml/100 ml deionized water). University Animal Care staff performed daily checks on animal welfare with Miller lab personnel providing oversight throughout the duration of the experiment.

### 2.3 Tetrode Implantation and Electrophysiology

#### Surgical procedure and implantation

Approximately two months post-virus injection, 17 finches, out of the 19 originally included (two birds died post-virus injection and prior to tetrode implantation), were anesthetized with isoflurane (2-3%, constant flow rate of 0.5-0.8 L of oxygen/min). While under anesthesia, they underwent acute electrophysiological recordings of right hemisphere Area X as a non-survival surgery approved under PI Miller’s IACUC Protocol #13-489. Breathing rate was monitored throughout the experiment, and an infrared heating pad (Kent Scientific) maintained body temperature at 37°C. The same stereotaxic coordinates were used for the AAV injections targeting Area X.

##### Ground wire insertion

A ground/reference wire was inserted into the left hemisphere, contralateral to the virus injection site (mid-sagittal sinus: 3.1 mm rostral-caudal, 1.62 mm medio-lateral). After performing the craniotomy, the dura was pierced with a 30G needle, and the end of the ground wire was lowered to a depth of 2-3 mm. Gel superglue was used to secure the wire to the acid-etched skull (acid from the Metabond kit). Zip Kicker (Pacer Technology) was used to rapidly cure the superglue.

##### Electrode array insertion

The existing craniotomy on the right hemisphere (used for virus injection) was expanded to a diameter of ∼1.5 mm to better accommodate the electrode array and to have access to dura that was undamaged during virus injection. Tetrode position was confirmed in Area X post-mortem following electrophysiological recordings (**Fig 2**).

**Fig 2.**
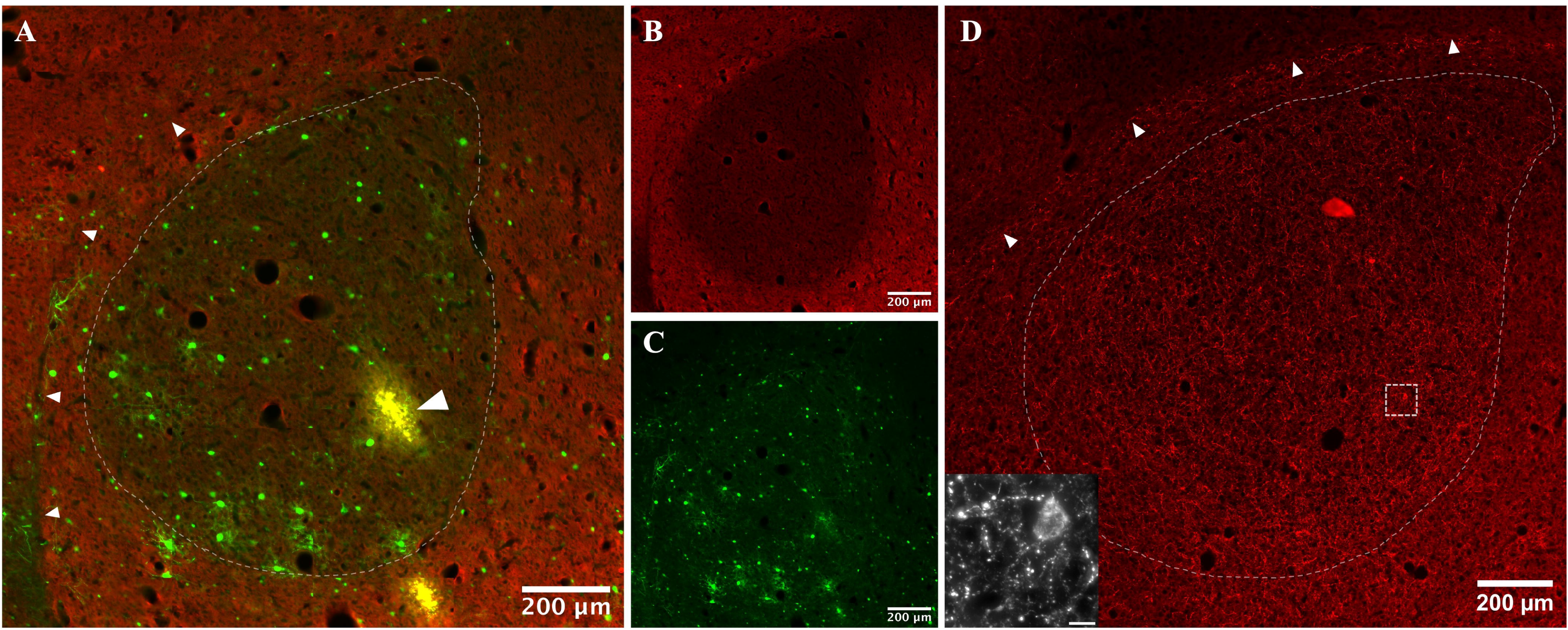
AAV expression in finch song nucleus Area. **X.** The Area X region is represented by the dotted line. Small arrowheads denote the border between the striatum and the nidopallium. Staining for alpha-synuclein (a-syn) protein is shown in red. **A)** An AAV control expressing finch has GFP labeled cell bodies (green) in Area X, and the large arrowhead denotes the last recorded tetrode position (yellow). **B)** A-syn protein expression is low in Area X compared to surrounding basal ganglia in this AAV-GFP finch. **C)** View of (A) showing the GFP channel only. **D)** An AAV-h*SNCA* expressing finch shows widespread expression of a-syn in processes throughout Area X. Inset in bottom left corner shows a zoomed in view of bead-like a-syn protein in the processes (inset scale bar = 10 µm).

#### Electrophysiological recordings

Neurons from Area X were recorded using a custom-made 800 µm diameter tetrode array composed of 8 twisted-wire tetrodes. Tetrodes were constructed from polyamide-insulated nickel chromium steel wire (∼12 µm in diameter; see **Fig 1C**). Prior to insertion, tetrodes were coated with DiI to allow anatomical tracing of the electrode path (**Fig 2A**). The tetrode array was lowered in ∼100 µm increments until the tip reached Area X at ∼3100 µm (right hemisphere) and allowed to stabilize for 5-10 minutes prior to recording. Neural data was acquired at 20 kHz using the INTAN neural recording system (Intan Technologies, Inc.). Neural recordings from Area X neurons occurred in 5-minute blocks. After each 5-min block of data was acquired, the electrode array was lowered ∼50 µm and another recording block was acquired. All recordings were restricted to Area X (3100-4100 µm). Data was spike sorted using Kilosort v2.5 (https://zenodo.org/records/4482749) (71) and Phy (https://github.com/cortex-lab/phy). Methods for cell-type estimation are described in Results.

#### Data/Statistical Analysis

All single-unit data was analyzed using Matlab (R2025a). All Matlab analysis code is available upon publication at: https://github.com/CowenLab/ZebraFinch. Out of the 17 finches implanted with the tetrode array, four finches in the ASYN group yielded no recordings (e.g. neurons were quiet), and one GFP control finch showed no virus expression detected in Area X post-mortem. These five finches were therefore excluded from our data analyses but humanely euthanized with the other finches at the end of that day’s electrophysiological recording period. Thus, n=12 finches (n=6/per AAV group) were used for electrophysiological and song analyse presented in the Results section.

##### Measurement of burst activity

The temporal structure of the firing activity of individual neurons was quantified using “local variance” (LV), a measure that assesses the variability of inter-spike intervals (ISIs) and is robust to changes in firing rate (72). LV is computed from the vector of ISIs for each neuron as shown in the formula below:

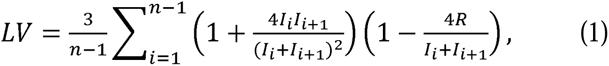

where *I_i_* and I*_i_*_+1_ are the *i*-th and *i* + 1 ISI and *n* is the number of ISIs. LV scores ∼0 indicate tonic firing, scores ∼1 show Poisson firing, and scores >1 indicate burst-like firing patterns.

Parametric and non-parametric tests were used for hypothesis testing. For group-level analysis of firing-rate data (n = 201 categorized neurons from n = 12 finches) non-parametric tests were used as firing rates do not follow normal distributions (73). Consequently, rank-sum tests were used for two-group comparisons and the Kruskal-Wallis test was used for multi-group data. The Holm-Bonferroni correction was used to correct for multiple comparisons. For normally distributed data, ANOVA and t-tests were used followed by Tukey or Holm-Bonferroni post-hoc corrections. Alpha was set to 0.05. Relationships between neural data and each bird’s song features (e.g., mean Entropy, Duration, and Amplitude) were investigated using a generalized linear regression model where the outcome variable was neural activity (e.g., firing rate) and the independent variables were song features and group category (ASYN or GFP) as described in **Results and S2 Fig**. Multicollinearity between independent variables can impair the ability to interpret regression models. We observed that some song features did indeed covary (r > 0.4 for 5 song feature pairs). To address this, we repeated the regression using the first three principal components of the independent variable matrix and compared results to those from the original regression model. As reported in **S2 Fig**, similar results were obtained from the original and dimensionality-reduced model.

### 2.4 Tissue processing and immunohistochemistry

Immediately following the end of the neural recording session, finches were humanely euthanized by isoflurane inhalant overdose and transcardially perfused with warmed saline followed by chilled 4% paraformaldehyde in Dulbecco’s Phosphate Buffer Saline. Fixed brains were cryoprotected in 20% sucrose overnight then cryosectioned in the coronal plane at 30 µm on a Microtome cryostat through Area X. Tissue was processed, following our previously published methods (45, 46) and summarized here: Hydrophobic borders were drawn on the slides, using a pap pen (ImmEdge, Vector Labs) followed by 3 × 5 minute washes in TBS with 0.3% Triton X (Tx). To block non-specific antibody binding, the tissue was then incubated for one hour at room temperature with 5% goat serum (Sigma #G-9023) in TBS/0.3% Tx then 3 x 5-minute washes in 1% goat serum in TBS/0.3% Tx were performed. A primary antibody was used against a-syn (1:250, rabbit, Proteintech 10842-1-AP, RRID: AB_2192672). GFP expression was robust enough to be visualized without an antibody.

The primary antibody was incubated in a solution of 1% goat serum in TBS/0.3% Tx overnight at 4°C. One tissue section per slide was used as a ‘no primary’ control and a pre-absorption control experiment was performed to validate antibody specificity in a companion study evaluating the distribution of a-syn protein following AAV injection (46). The next day, sections were washed 5 x 5 minutes in TBS/0.3% Tx and incubated for three hrs at room temperature in a fluorescently conjugated secondary antibody (1:1000: goat anti-rabbit 647 A-21245, RRID:AB_2535813). After secondary incubation, sections were washed 3 x 10 minutes in TBS followed by 2 x 5 washes in filtered TBS. Slides were then cover-slipped in Pro-Long Anti-Fade Gold mounting medium (Molecular Probes, P36930). Tissue was imaged using a Zeiss Axio Observer 7 with Apotome III Microscope under the University of Arizona Imaging Cores - Optical Core Facility (RRID:SCR_023355).

## Results

### Virally-driven expression is confirmed in song nucleus Area X

**Fig 2** is a representative example of AAV expression in Area X. Two months post-virus injection, we detected GFP cell bodies in Area X (**Fig 2A,C**) accompanied by low a-syn protein expression (**Fig 2B**). By contrast, a finch overexpressing a-syn showed staining in processes and cell bodies (**Fig 1D**). These results are consistent with our two prior publications (45, 46).

### Extracellular identification of neuronal subtypes in anesthetized finches

Area X neurons share much in common with neurons in the mammalian striatum and pallidum. For example, the most abundant Area X neuron type is homologous to the mammalian medium spiny neuron (MSN,(58)), with Area X and mammalian MSNs sharing wide waveforms, low firing rates, and bursting activity (74–76). There is also evidence that Area X contains striatal fast-firing interneurons, cholinergic interneurons, and low-threshold spiking neurons (48, 74, 77). Furthermore, Area X contains two classes of fast-firing pallidal-like neurons: local-circuit (HF-1) and thalamus-projecting (HF-2) (59, 78). These six cell classes were identified by their firing rates and waveform features using multiple approaches from this literature, including correlating neuronal activity *in vivo* with birdsong and patch-clamp recording in brain slices. Our present study is constrained by the fact that all data was acquired from extracellular electrodes implanted in anesthetized finches. We found no prior publications with recordings from Area X of adult anesthetized finches; prior work includes some anesthetized recordings from lMAN and Area X in the context of juvenile song learning (79). Consequently, our classification was informed by criteria developed from *in vivo* recordings from Area X in singing finches and restricted to waveform shape, firing rate, and assessment of bursting activity in the following five identified categories (74, 78, 80).

Neuron categorization and criteria (results shown in **Fig 3**):

1. **Wide-Low-Rate (WLR) Criteria**: Neurons firing ≤ 6 Hz with a peak half width > 0.25 ms or a peak-to-trough width of > 0.35 ms (**Fig 3 A,D,E**). WLR neurons were also more ‘bursty’’ relative to other neuronal types (**Fig 3F**) as measured by Local Variance (LV, see Methods (81)) and were the most abundant of the five identified cell types (**Fig 3C**). Together, these features suggest that WLR cells were MSN-like given evidence that MSNs are the most prominent cell type in the striatum, exhibit wide waveforms, have low firing rates, and express bursting activity (74) (75, 76).
2. **Wide-High-Rate (WHR) Criteria**: Neurons with a firing rate between 20 and 50 Hz with a peak half width > 0.25 ms or a peak-to-trough width of > 0.35 ms (**Fig 3A,D,E**). Post-classification analysis also showed that these neurons fired tonically (LV < 1, see **Fig 3F**). Tonically active WHR cells have features consistent with striatal cholinergic interneurons (74).
3. **Narrow-Low-Rate (NLR) Criteria**: Narrow waveform neurons firing ≤ 6 Hz (74) having a peak half width < 0.25 ms or a peak-to-trough width of < 0.35 ms (**Fig 3A,D,E**).
4. **Narrow-High-Rate (NHR) Criteria**: Narrow waveform neurons with rates ≥ 8 Hz but < 50Hz (**Fig 3A,D,E**). Narrow waveforms are characteristic of local circuit fast-firing inhibitory neurons.
5. **Very-High-Rate (VHR) Criteria**: Neurons firing ≥ 50 Hz (**Fig 3A,D,E**). Area X pallidal-like projection neurons are characterized by exceedingly high firing rates (74, 78). Consequently, we established a category for neurons with rates ≥ 50 Hz. Waveform features were not used to further subdivide neurons in this group given its small size (n = 18 neurons, **Fig 3C**). Analysis of VHR firing activity revealed that these neurons exhibited decidedly tonic activity, with a mean LV of 0.51 (**Fig 3F**). These data suggest that the identified VHR neurons are type-2 pallidal-like projection neurons (HF-2 in (78) (82)) given the presence of tonic firing and relatively wide waveforms. Furthermore, the small size of this group is consistent with other reports that suggest PAL neurons comprise approximately just 5.4% of the Area X cell population (58).

**Fig 3.**
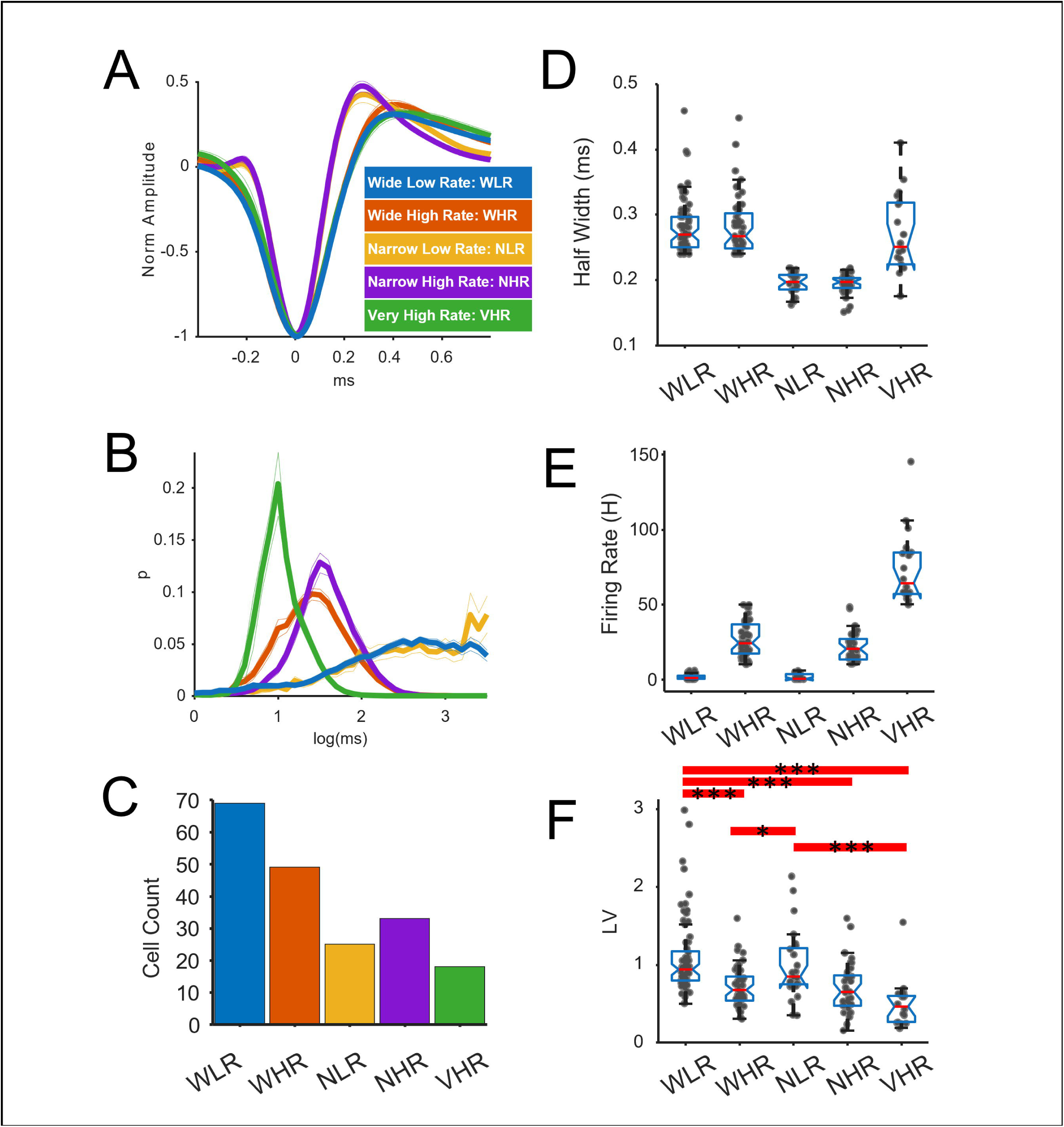
Identification of five classes of Area X neurons in anesthetized finches. **A)** Mean shape for each of the five identified classes of neurons (n = 201 neurons from n = 12 finches). Neurons were categorized based on spike width and firing rate (see Results). Amplitude is normalized such that the minimum (trough) = −1. Colors indicate neuron subtype (see key). **B)** Inter-spike interval distributions for each neuronal subtype. Y-axis indicates probability (p). **C)** Histogram indicating the number of identified neurons in each category. **D)** Box plot of the half-width (ms-milliseconds) for each neuronal group. Each dot indicates an individual neuron. Horizontal red lines indicate the median. Hypothesis tests were not performed as half-width was a key criterion for classification. **E)** Firing rates for each neuronal group (H-Hertz). Each dot indicates an individual neuron. Hypothesis tests were not performed as firing rate was a key criterion for classification. **F)** Burst firing differed between groups. While neurons were not categorized based on burst-firing, the measure of bursting (Local Variance-LV) indicated that Wide-Low-Rate neurons had increased burst firing (LV>1) relative to other groups (ANOVA p < 0.000001, Tukey post-hoc comparison. Thick red lines indicate significant post-hoc differences (*p < 0.05, **p < 0.01 (not shown), ***p < 0.001). The red lines in the box plots indicate the median and the whiskers indicate 1.5X the Interquartile Range.

### Reduced Area X firing rates in ASYN overexpressing finches

We predicted that a-syn expression would reduce putative MSN (WLR) neuron activity, thus releasing putative PAL neurons (VHR) from inhibition. Consequently, we investigated whether firing rates of WLR neurons were reduced in the ASYN group and whether rates in VHR neurons were increased. Visual inspection of raster plots of WLR activity (**Fig 4A-B**) suggested reduced WLR firing activity in the ASYN group. Statistical comparison of firing rates of GFP and ASYN neurons identified a significant difference (**Fig 4C**, rank-sum, p = 0.005 before correction and p = 0.026 after Holm-Bonferroni correction). No significant between-group differences were identified in the other neuronal classes, including the VHR neurons.

**Fig 4.**
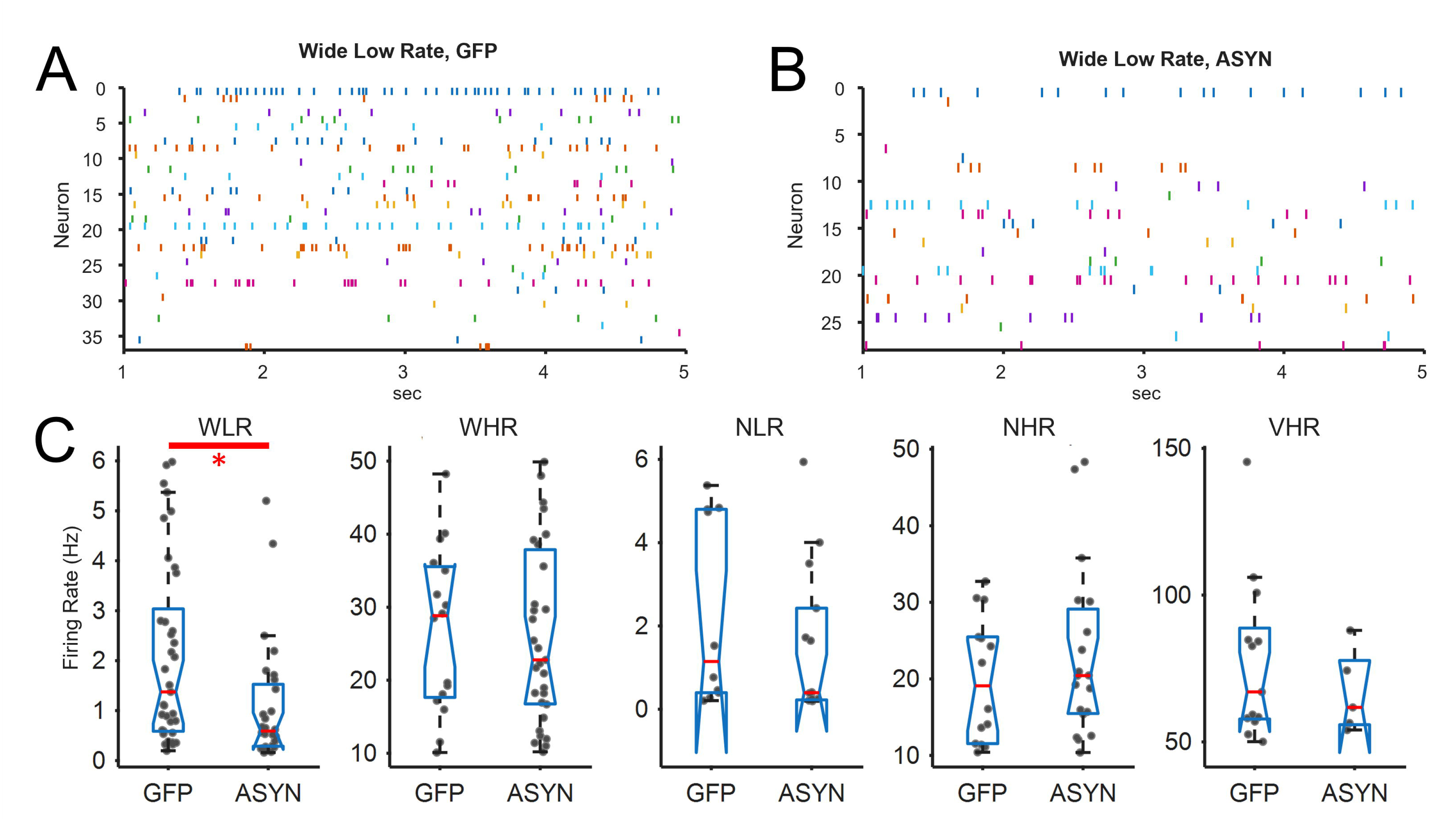
Reduced firing rates of WLR neurons in the ASYN group. **A-B)** Raster plots of WLR in the GFP and AAV-h*SNCA* (ASYN) groups. The x-axis shows time in seconds. Each row is a raster for a distinct neuron. **C)** Between-group comparison (GFP vs. ASYN) for each of the five neuron types. WLR neurons were only found in 4/6 finches. Firing rates for WLR neurons were lower in the ASYN group relative to the GFP control (rank-sum test, p = 0.005 prior to correction, *p = 0.026 after Holm-Bonferroni correction).

### Burst firing was not changed in ASYN overexpressing finches

While burst firing is a canonical property of MSNs, MSN burst firing may become dysregulated in PD. Indeed, altered MSN burst firing has been reported in the 6-OHDA mouse model of PD (83), non-human MPTP primate models (84), and in human PD patients (64). Consequently, we predicted that burst firing will be affected by ASYN overexpression in finch Area X. To investigate this, the local variance (LV) measure (72), see Methods) was used to evaluate the pattern of inter-spike intervals and quantify the degree that neuronal activity was either tonic (low LV), Poisson (LV near 1), or bursty (LV > 1). Contrary to our prediction, analysis of spiking statistics with LV did not identify any difference between GFP and ASYN groups in LV values for any neuronal subtype (**Fig 5**, t-test, all p > 0.05 after Holm-Bonferroni post-hoc correction). Thus, while ASYN significantly reduced firing rates of WLR neurons, these changes were not accompanied by a change in burst-firing.

**Fig 5.**
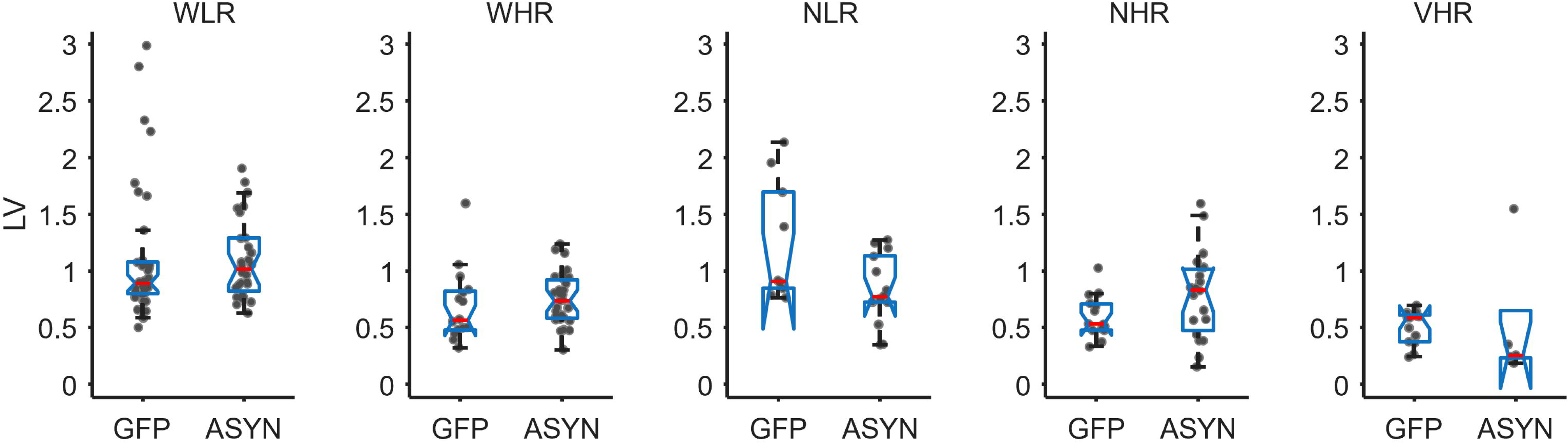
Local Variance (LV), a measure of bursting, is not different between ASYN and GFP control groups. Between-group comparisons (GFP vs. ASYN) were made for each of the five neuron types, and no between-group differences were identified in any of the neuron classes (rank-sum test, all p > 0.05 after Holm-Bonferroni post-hoc correction).

### Extracellular waveform shape was altered in wide-waveform neurons

While waveform features and firing rates are commonly used to classify neurons into putative subtypes (74, 80, 85), inferring physiological properties of neurons from extracellular recordings is highly problematic as waveform features vary as a function of recording location relative to the morphology of the neuron (86–88). Even so, we did observe consistent differences in extracellular waveform shapes between ASYN and GFP animals that encourage future investigations. Specifically, we found that WLR and WHR neurons showed a large positive extracellular ‘rebound’ following the initial trough (**Fig 6A**). This was determined by first normalizing the waveform of each neuron such that the trough = −1 and baseline was zero. Consequently, positive values indicate proportional changes above baseline. The “Rebound Peak” for each neuron was measured as the normalized amplitude of the largest peak following the time of the trough (t = 0 ms). Values for each neuron type are shown in **Fig 6B** for the GFP and ASYN groups. Rank-sum tests were performed, and significant between-group differences were identified in the WLR and WHR groups (p_WLR_ = 0.003, p_WHR_ = 0.0002 after Bonferroni-Holm correction).

**Fig 6.**
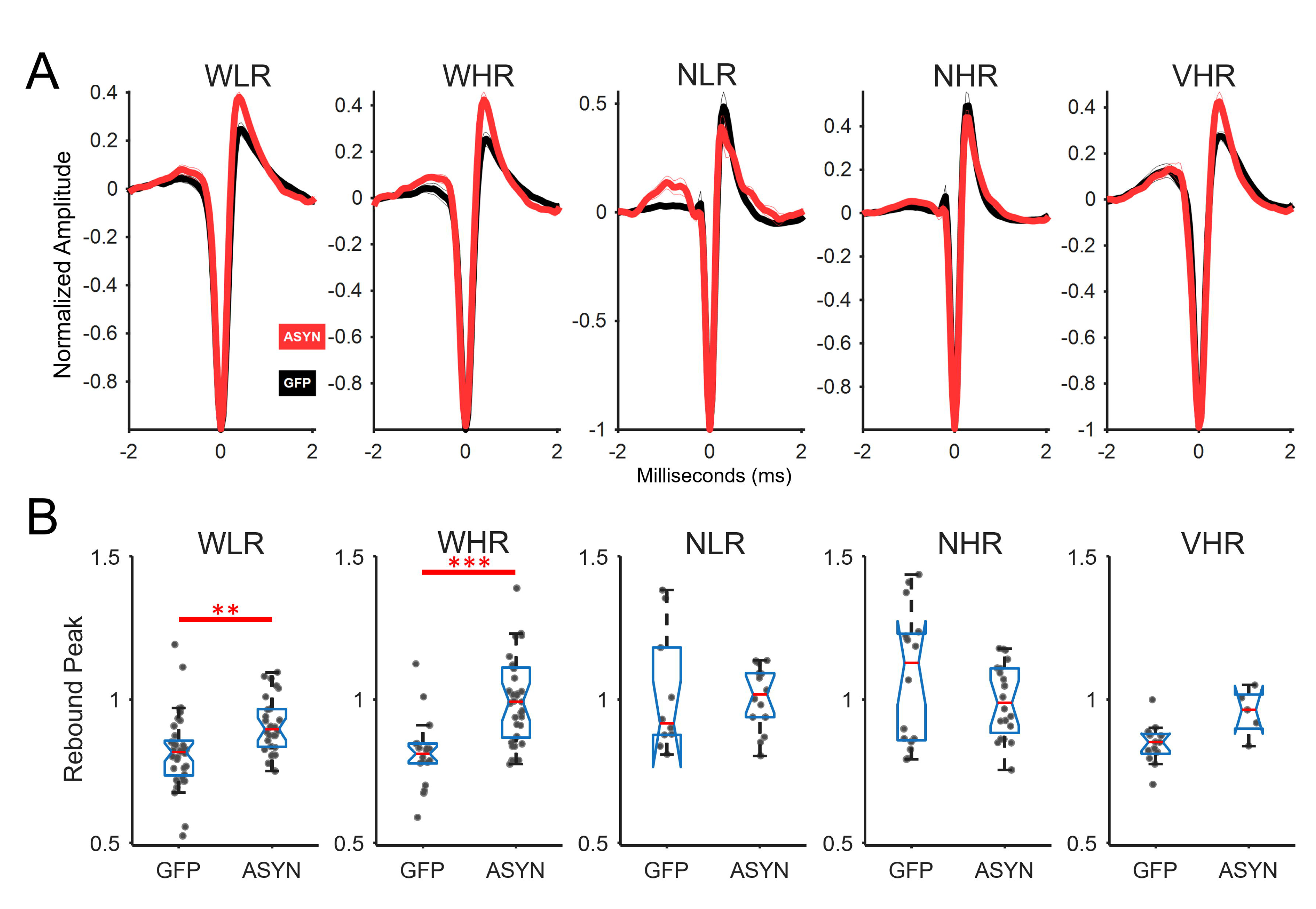
ASYN group shows enhanced post-peak rebound for wide-waveform neurons. **A)** Mean waveforms for the ASYN and GFP groups for each neuron type. For each waveform, data was normalized such that the distance from baseline to the trough = −1. Inspection of these plots suggested that wide-waveform (WLR, WHR) ASYN neurons had a larger rebound following the initial trough. **B)** Statistical analysis of the rebound was performed by measuring the normalized amplitude of the largest peak following t=0 (rebound peak) for each neuron (dots). These values were compared between groups using rank-sum tests and p values were adjusted using the Holm-Bonferroni correction. **p<0.01, ***p<0.001. Error bars in box plots indicate ±SEM.

### No observed correlation between song features and neuronal firing properties

Next, we investigated whether ASYN and GFP birds differed in their song features and if variation in mean WLR neuronal activity in each bird correlated with song features. Analysis of syllable-level features failed to identify any between-group difference (Mann-Whitney U test, p > 0.05, **S1 Fig, S1 Table**).

Investigation of whether firing rates correlated with song features was performed by first calculating the mean firing rate of WLR neurons (the largest group and the putative MSNs) for each bird. Two birds had to be excluded from the analysis as no WLR neurons were identified in these animals. Mean WLR firing rates were used as the outcome variable for a generalized linear model where the independent variables were song features (mean amplitude, duration, and entropy and CV amplitude, duration, and entropy) and the group (ASYN or GFP). No significant relationship between song features and firing rate was identified (F(10,2) = 11.7, p = 0.082);**S2 Fig**.

## 4. Discussion

The goal of this study was to obtain new information about the cellular and network-level activity affected by overexpression of human a-syn protein in an adult male zebra finch model. To carry out this goal, we performed extracellular recordings from song nucleus Area X of the basal ganglia in anesthetized adult male finches and identified five distinct cell types. Our main finding was that wide, low firing rate (WLR) neurons, putative MSNs, show reduced firing activity and enhanced post-inhibitory rebound in the a-syn overexpression condition compared to controls. This aligns with our working hypothesis that abnormally elevated a-syn protein levels in Area X will decrease MSNs firing rate. We also predicted that GP-projecting neurons would show reduced firing in Area X. VHR neurons may represent these GP-projecting neurons, but they did not show a difference in firing rate between the ASYN and GFP control groups. This finding may reflect low statistical power given the small number of VHR neurons recorded.

A benefit of anesthetized recordings is that they allow for high data throughput, enabling the recording of a substantial proportion of neurons without confounds due to animal movement. A limitation of this approach in our finch model is that on-going real time modification of neural activity related to song production cannot be evaluated. It is also likely that neuronal activity was dampened by isoflurane anesthesia. While information is scarce on the effects of isoflurane on *in vivo* recordings from the striatum of naïve finches or rodents, patch clamp studies in rat brain slices show that brief isoflurane exposure can dampen sodium currents in hippocampal neurons (89) and reduce spiking and bursting modes in thalamic neurons (90). Our future investigations will compare isoflurane-induced anesthesia on Area X of naïve finches versus awake, singing finches where we can examine how a-syn modulates neuronal dynamics under more physiologically relevant conditions.

The observed reduction in WLR (putative MSN) firing rates observed in Area X of the ASYN finches and lack of bursting activity contrasts with some measures of MSN activity reported in human PD patients and primate PD models. For example, striatal projection neurons in the human basal ganglia of PD patients undergoing deep brain stimulation had high frequency neuronal firing patterns resembling the advanced parkinsonism of the Non-Human Primate model, with hyperactivity and “bursty” neuronal firing activity (14, 64). In a mouse model of bilateral dopamine depletion (via the neurotoxin 6-OHDA) in the dorsolateral striatum, single unit recordings from MSNs showed reduced bursting activity during movement. Intriguingly, they classified MSNs into two types with type 1 showing increased firing during movement and type 2, decreased firing with no difference in firing rate when the mouse was stationary (83). In freely moving rats, 6-OHDA treatment increased oscillatory synchronization in the striatum (91). Future recordings in singing, ambulatory finches with the ASYN phenotype will determine if hyperactivity of MSNs and differential sub-types exist, evidence for dopamine loss, and how these various sub-components contribute to changes in real-time song output.

The reduced firing rates of WLR neurons in ASYN expressing finches may reflect reduced mean frequency of spontaneous excitatory postsynaptic glutamatergic currents from cortical song nucleus lMAN onto Area X MSNs and GP neurons. Striatal recordings from h*SNCA* overexpressing mice show reduced glu currents (12). Future work using electrophysiological recordings from Area X brain slices *in vitro* can determine whether glu input is perturbed in our ASYN condition.

ASYN-driven abnormal changes in synaptic input to Area X may also be accompanied by altered intrinsic membrane properties. Both the WLR and WHR neuronal classes showed greater positive rebound potential following the initial spike in the ASYN expressing group compared to GFP controls. While extracellular recordings of action potentials cannot reliably identify after-hyperpolarization (AHP) currents, the observed rebound potential occurs during the typical time interval of the AHP, suggesting that membrane properties in the ASYN neurons are altered. The AHP is regulated by K^+^ currents, and several classes of K^+^ channels are implicated in altered neuronal excitability in PD models (92). The significantly larger rebound potential in the ASYN expressing group may reflect an enhanced AHP-related conductance. Patch-clamp electrophysiology in brain slices is a necessary approach to confirm true fluctuations in membrane properties during the AHP period. Changes in AHP-related conductance could result from increased SK/BK channel activity (93) (94)) or a reduction in I_h_ current (95, 96), leading to greater hyperpolarization following spikes and dampened excitability. In nigral DA neurons, abnormal a-syn oligo aggregates increase the conductance of K^+^_ATP_ channels leading to reduced firing (97). In our finch model, a-syn overexpression could progressively decrease intrinsic excitability in MSNs via these channels, and/or others. To evaluate the specific role of a-syn expression on MSNs, future directions will employ a custom-made AAV containing a CAMKII promoter that has been previously used in naïve finches to evaluate MSNs firing activity (98). By selectively overexpressing a-syn in MSNs, we can evaluate the relationship between the extent of protein pathology and the neural activity changes underlying song. In a recent publication, we found that regional changes in virally-expressed human a-syn protein within Area X correlated with changes in syllable level features. For example, we detected a positive correlation between right hemisphere a-syn proteinopathy and a reduction in the variation of harmonic syllable duration, among other features, correlated to the right hemisphere Area X (46). Our neural recordings here are also from right hemisphere Area X; the contribution of the left Area X is not known. Prior studies suggest a transition in the unilateral activation of the hemispheres: right-side dominance of song nucleus HVC is detected in adult birdsong (99) while the left-side dominates during juvenile learning (100). Thus, when assessing cell-type changes in electrical activity within brain nuclei aligned with real-time singing behavior, factors to take into account include the recording hemisphere, the distribution of protein expression, and the presence of pathology, e.g. a-syn aggregates in individual cell types.

## Conclusions

Our study takes an important first step forward in evaluating the consequences of a-syn overexpression on the activities of neuronal subtypes involved in vocal behavior. A growing body of research has elucidated the role that neurons in the cortico-striatal-thalamic circuit play in male zebra finch song production. By introducing human a-syn to the zebra finch model, this study supports a systems- and cellular-level investigation of human vocal deficits in PD. This is important given that current medications and brain stimulation used in the treatment of PD fail to resolve the vocal deficits and can worsen them (101–103). Data from pre-clinical animal models such as the songbird can provide important mechanistic insight needed for the development of new brain-targeted cellular therapies for the vocal dysfunction.

## CRediT Contributions

### CRediT authorship contribution statement

Brian R. Dominguez: Data curation, Investigation, Methodology, Validation, Visualization, Writing-Original Draft, Writing-review & editing. Gabriel Holguin: Data curation, Formal Analysis, Investigation, Methodology, Software, Validation, Visualization, Writing-Original Draft, Writing-review & editing. Madeleine S. Daly: Data curation, Formal Analysis, Investigation, Methodology, Validation, Writing-review & editing. Reed T. Bjork: Investigation, Methodology, Validation, Visualization, Writing-review & editing. Stephen Cowen: Conceptualization, Data Curation, Formal Analysis, Investigation, Methodology, Software, Supervision, Validation, Visualization, Writing-Original Draft, Writing-review & editing. Julie E. Miller: Conceptualization, Data curation, Formal Analysis, Funding acquisition, Investigation, Methodology, Project administration, Resources, Supervision, Validation, Visualization, Writing-original draft, Writing-review & editing.

## Acknowledgements

We thank Kathy Dillon, Michelle Gordon, and Mauricio Serna for technical contributions along with University of Arizona Animal Care staff and Mr. Douglas Cromey of the University of Arizona Imaging Cores - Optical Core Facility (RRID:SCR_023355).

## Supporting Information

**S1 Table 1. Song data and statistics.** For further information, see legends embedded into the tabs of this excel file.

**S1 Fig. Exemplar song motif and group level comparisons of song syllable data.**

**A)** Spectrogram of song motif from a pre-injected GFP control bird (as in **Fig 2A-C**) with time in seconds on the x-axis and frequency in kilo (k) Hertz on the y-axis. Unique syllables are assigned a letter (A-E) and classified into harmonic, noisy, or mixed syllables (68). **B-E)** The mean and SEM scores shown for syllable level acoustic features (duration, amplitude, and entropy) for the ASYN (red) and GFP (black) control groups. Raw scores can be found in the **S1 Table**. The y-axis is the normalized score calculated from dividing the post-AAV injection scores by the pre-injection scores. Each dot represents an average score per finch. **B)** Comparisons of all syllable types combined or by individual syllable types **(C-E)** reveal no group differences (Mann Whitney U, p>0.05). Individual variation in CV scores is detected.

**S2 Fig. No significant correlation between song features and firing rate.** Scatter plots showing the mean firing rate of WLR neurons averaged for each animal (dot) organized by GFP (black) and ASYN (red) groups. Plots were generated for each song feature, representing all syllable types combined. Generalized linear regression (see Results) did not identify a relationship between firing rate and song features or group (p = 0.0815).

## Notes

### Competing Interest Statement

I have read the journal's policy and the authors of this manuscript have the following competing interests: J.E. Miller is a 2025 member of the T2 Targets to Therapies advisory working group for the Michael J. Fox Foundation for Parkinson?s Research.

### Summary of Updates

The main manuscript file text has been updated. The Supplemental material is also now provided in the form of a word document containing Supplemental Figures 1-2 and an excel file, Supplemental Table 1, containing the song data and statistics. All of the main figures are re-included here in this version.

