## Supplemental Figures 1-2 for "Alpha-synuclein overexpression reduces neural activity within a basal ganglia vocal nucleus in a zebra finch model"

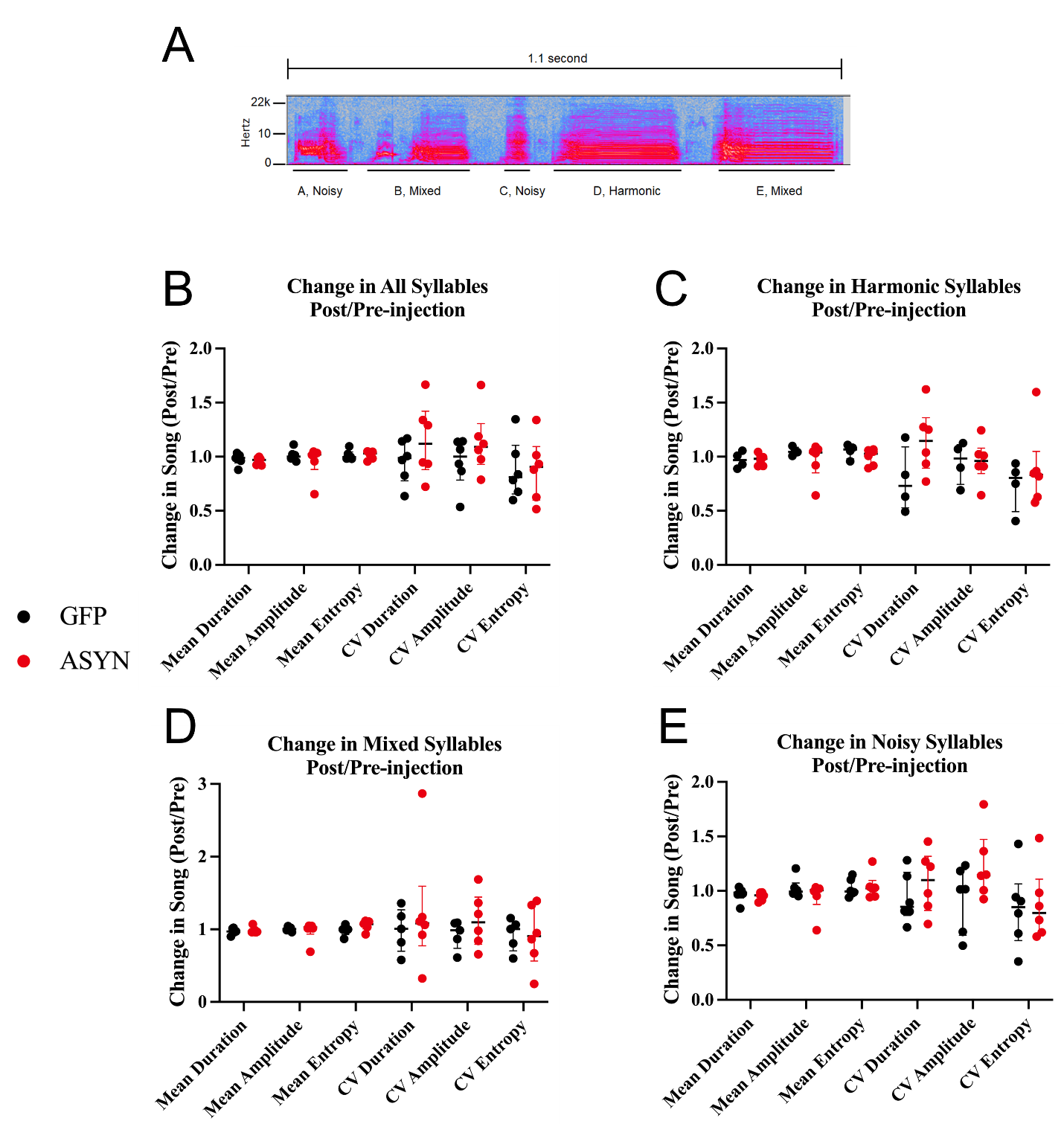

**S1 Fig. Exemplar song motif and group level comparisons of song syllable data.** **A)** Spectrogram of song motif from a pre-injected GFP control bird (as in **Fig 2A-C**) with time in seconds on the x-axis and frequency in kilo (k) Hertz on the y-axis. Unique syllables are assigned a letter (A-E) and classified into harmonic, noisy, or mixed syllables {Badwal, 2018 #4296}. **B-E)** The mean and SEM scores shown for syllable level acoustic features (duration, amplitude, and entropy) for the ASYN (red) and GFP (black) control groups. Raw scores can be found in the **S1 Table**. The y-axis is the normalized score calculated from dividing the post-AAV injection scores by the pre-injection scores. Each dot represents an average score per finch. **B)** Comparisons of all syllable types combined or by individual syllable types **(C-E)** reveal no group differences (Mann Whitney U, p>0.05). Individual variation in CV scores is detected.

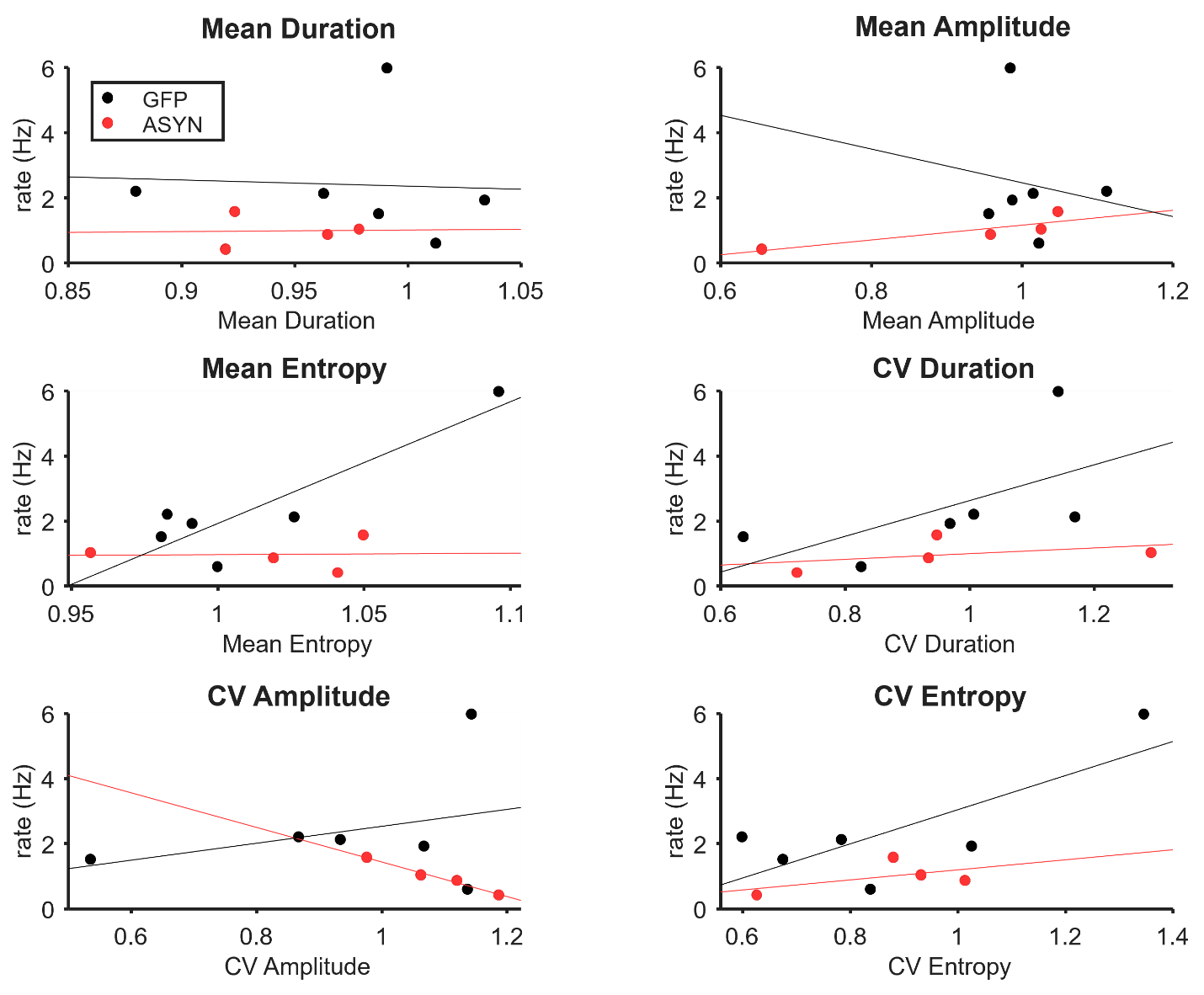

**S2 Fig.** **No significant correlation between song features and firing rate.** Scatter plots showing the mean firing rate of WLR neurons averaged for each animal (dot) organized by GFP (black) and ASYN (red) groups. Plots were generated for each song feature, representing all syllable types combined. Generalized linear regression (see Results) did not identify a relationship between firing rate and song features or group (p = 0.0815).
